# Twinfilin is a potent uncapper of actin capping protein and modulates actomyosin contractility in the *C. elegans* spermatheca

**DOI:** 10.1101/2025.06.17.660173

**Authors:** Anupreet Saini, Shir Kreizman, Ekram Towsif, Jonathan Martinez-Lopez, Iska Maimon Zielonka, Anat Nitzan, Lee Rudnik, Shashank Shekhar, Ronen Zaidel-Bar

## Abstract

The actin cytoskeleton is dynamically remodelled by conserved regulators to control cellular and tissue mechanics. While the functions of these proteins are well studied, how they drive tissue-specific contractility remains unclear. Twinfilin, an actin uncapper and depolymerase, has not previously been linked to tissue contractility. Here, we show that the sole twinfilin ortholog in *C. elegans*, TWF-2, regulates actomyosin contractility in the spermatheca. TWF-2 localizes to the spermathecal cortex via interactions with α-spectrin (SPC-1) and β-spectrin (UNC-70). *In vitro*, TWF-2 promotes barbed-end depolymerization and rapidly removes CAP-1 from actin filaments. *In vivo*, embryonic lethality caused by CAP-1 depletion is partially rescued by simultaneous loss of TWF-2. Similarly, loss of the contractility regulator SPV-1 leads to elevated F-actin and phosphorylated myosin, causing hypercontractility. Notably, removing TWF-2 suppresses this hypercontractility by reducing F-actin levels— without affecting myosin or its phosphorylation—highlighting a specific role in F-actin regulation. Together, these findings show that TWF-2 modulates actin dynamics in a tissue-specific manner. This work provides the first *in vivo* evidence that twinfilin regulates contractility, and reveals how its interactions with capping protein and spectrins help maintain balanced actomyosin levels in the spermatheca.

## Introduction

The actin cytoskeleton is a dynamic filamentous protein network essential for fundamental cellular processes such as cytokinesis, motility and cell shape changes. Actin-binding proteins, including those that bind preferentially to actin monomers or to specific filament ends, play a critical role in regulating the actin cytoskeleton’s architecture and dynamics, to facilitate execution of these processes at the right time in the right place. Among these proteins, twinfilin is an evolutionary conserved key regulator of actin dynamics, with diverse roles across species (Ulrichs & Shekhar, 2025)

In yeast, twinfilin deletion causes morphological defects in budding cells (Goode et al., 1998). In Drosophila, twinfilin regulates bristle formation (Wahlström et al., 2001), border cell migration and axonal outgrowth (D. Wang et al., 2010). In mammals, while initially reported to be dispensable for mouse development (Nevalainen et al., 2011), twinfilin is critical for platelet function in mice (Stritt et al., 2017) and neurite growth and spine density in cultured rat cells (Y.-J. Wang et al., 2021; Yamada et al., 2007). In humans, both reduction and overexpression of twinfilin have been implicated in cancer progression (Bockhorn et al., 2013; Samaeekia et al., 2017; Zhai et al., 2023).

Despite species-specific phenotypic outcomes, twinfilin universally interacts with actin to modulate its dynamics. Initially established as an actin monomer binding protein (Goode et al., 1998b; Palmgren et al., 2001a; Vartiainen et al., 2003), it is now known that twinfilin interacts with the barbed end of F-actin, in a context dependent manner, to accelerate the depolymerization of filaments (Johnston et al., 2015; Moseley et al., 2006; Shekhar et al., 2020) or to cap the actin filament’s barbed ends (Helfer et al., 2006). Central to twinfilin’s function is its relationship with actin capping protein (CP), a major barbed end binding protein that terminates filament elongation (Falck et al., 2004; Palmgren et al., 2001b).

Twinfilin directly binds capping protein via its C-terminal tail which contains a CPI motif. Although it was initially thought to be a pro-CP factor owing to its protection of CP from CARMIL (Johnston et al., 2018), later studies showed that twinfilin functions as an uncapper that displaces CP from filament ends (Hakala et al., 2021; Mwangangi et al., 2021; Reddy et al., 2024; Ulrichs et al., 2023). Interestingly, although mammalian twinfilin on its own is a weak uncapper and only accelerates uncapping sixfold (Hakala et al., 2021), in the presence of formin mDia1, the two proteins synergize to accelerate uncapping by over 300-fold (Reddy et al., 2025). These findings highlight the complex regulation of actin dynamics by twinfilin, where its functional output is context-dependent (Ulrichs & Shekhar, 2025a). While their biochemical interplay is well-studied *in vitro* , the functional consequences of the interaction between twinfilin and actin CP in tissues remain poorly understood.

To address the role of these conserved actin regulators *in vivo*, we used *C. elegans* as a model system. Specifically, we focused on its reproductive system, which consists of two U-shaped gonads. Each gonad is comprised of somatic tissues, which house germ cells that differentiate into sperms and oocytes, with sperm stored in a specialised contractile fertilization chamber called spermatheca. CP is widely expressed in the *C. elegans* reproductive system, including the germline and spermatheca. Its role in germline architecture has been previously established, where its depletion leads to a hypercontractile germline through increased F-actin and myosin levels (Ray et al., 2023). The spermatheca has a highly organised actomyosin network which regulates its contractility over ∼150 ovulation events, making it an ideal system for dissecting the role of actin regulators in contractility. Its contractility is regulated by various proteins, including a RhoGAP called spermatheca physiology variant-1 (SPV-1), which acts as a mechanosensor and supresses contractility through inactivation of RHO-1 (Tan & Zaidel-Bar, 2015). Additionally, spectrins contribute to spermathecal contractility and maintain structural integrity of the actin cytoskeleton (Wirshing & Cram, 2018). However, no direct association between twinfilin and spectrin has been previously reported. Moreover, a role for twinfilin in regulating actomyosin contractility is still undescribed.

In this study, we characterize the expression, localization, and function of the only twinfilin in *C. elegans,* TWF-2. Specifically, we investigate how TWF-2 interacts with CP and spectrins, and how it modulates actomyosin contractility in the spermatheca. We demonstrate that TWF-2 is widely expressed in *C. elegans* and localizes to the spermathecal cortex in a spectrin-dependent manner. Furthermore, we show that the loss of TWF-2 mitigates the hypercontractility phenotypes caused by CP and SPV-1 dysfunction. *In vitro* assays confirm the direct association of *C. elegans* CP and TWF-2, providing mechanistic insight into their functional relationship. Together, these findings establish a role for twinfilin in tissue contractility and establish a framework for understanding its potential functions in other contractile tissues across species.

## RESULTS

### TWF-2 is widely expressed in contractile tissues and its cortical localization in the spermatheca is dependent on spectrins

To examine the expression pattern and subcellular localization of TWF-2 in adult *C. elegans* hermaphrodites, we tagged endogenous TWF-2 with the fluorescent marker GFPnovo (henceforth referred to as GFP) at its N-terminus (Figure 1A) using CRISPR/Cas9-mediated genome editing. To verify that tagging TWF-2 did not disrupt development, we measured the brood size and embryonic lethality of the GFP::TWF-2 strain and found them to be comparable to wild-type (Figure S1A, B).

**Figure 1:**
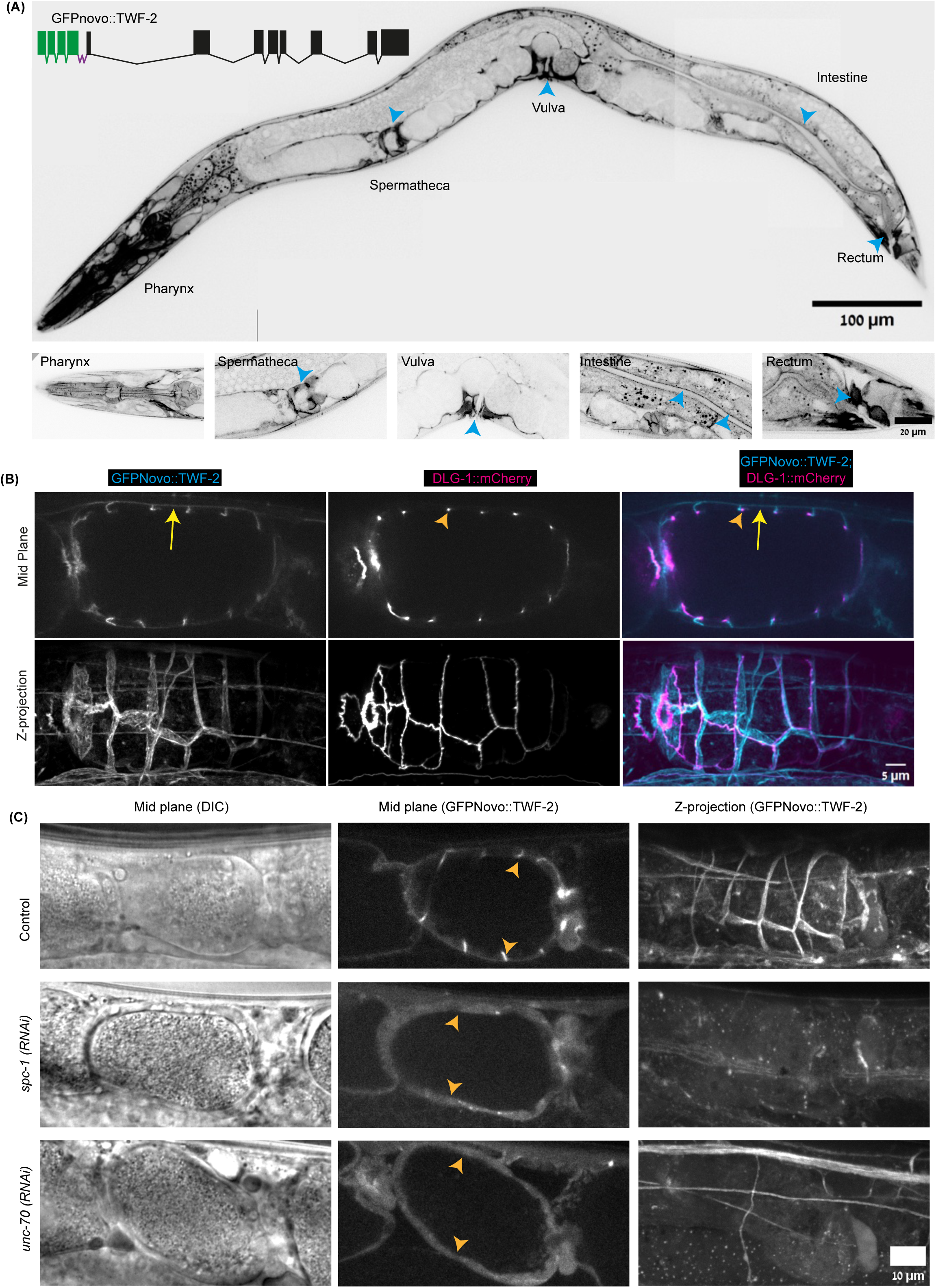
**Localization of endogenous GFP-tagged TWF-2 in *C. elegans* tissues** A. Endogenous GFP::TWF-2 expression pattern in adult hermaphrodites. Top: Representative mid-Z projections showing GFP::TWF-2 expression in contractile tissues (pharynx, spermatheca, vulva, intestine, and rectum; blue arrowheads). Schematic inset: Representation of the CRISPR design for GFPnovo::TWF-2, with exons (black boxes), introns (connecting black lines), GFPnovo (green) and linker (purple). Bottom: Magnified views of indicated regions are shown. B. Subcellular localization of GFP::TWF-2 (cyan) relative to the apical marker DLG-1::mCherry (magenta) in spermathecal cells. Single mid-plane (left) and maximum Z-projection (right) views demonstrate basolateral TWF-2 localization (yellow arrows) distinct from apical DLG-1 (orange arrowheads). C. RNAi mediated depletion of spectrins disrupts TWF-2 cortical localisation. Representative images of spermathecae from worms treated with control, *spc-1* and *unc-70* RNAi. Cortical GFP::TWF-2 signal (orange arrowheads) is disrupted upon spectrin knockdowns. N≥2 biological replicates with ≥40 worms analysed per condition.

Imaging of GFP::TWF-2 revealed widespread expression across multiple tissues, with particularly prominent expression in the pharynx, spermatheca, vulva, intestine, and rectum, which are all contractile tissues (Figure 1A). This spatial distribution suggested a potential role of TWF-2 in regulating non-muscle actomyosin contractility in *C. elegans*.

Next, we examined its subcellular localization and chose to focus on the spermatheca. Within spermathecal cells, we observed TWF-2 to be concentrated near the plasma membrane, in what appeared to be cortical localization. To validate its localization at the cellular cortex we crossed TWF-2::GFP with DLG-1::mCherry, a known apical junction protein, and observed that TWF-2 mainly localised to the basolateral membranes of spermathecae (Figure 1B). To identify proteins that interact with TWF-2 and possibly regulate its subcellular localisation, we performed co-ImmunoPrecipitation (co-IP) followed by Mass Spectrometry (MS) using the GFP::TWF-2 strain, with N2 as a control (Figure S1C). Compared to the N2 control, TWF-2 was enriched approximately 2000-fold in the GFP::TWF-2 sample, confirming the pull down assay. Based on enrichment scores and TWF-2’s known role as an actin-interacting protein, we shortlisted cytoskeletal candidates for further investigation (Table S1A). These candidates were depleted by RNAi followed by evaluation of TWF-2 localization in the GFP::TWF-2 strain.

From the candidate genes screened, RNAi knockdown of *spc-1* (α-spectrin) and *unc-70* (β-spectrin) specifically disrupted GFP::TWF-2 cortical localization in the spermatheca (Figure 1C ), and not in other contractile tissues where TWF-2 is expressed (Figure S1D), suggesting a tissue-specific role of spectrins in the cortical localization of TWF-2 in the spermatheca. Notably, *twf-2* RNAi, which knocked down 76% of TWF-2, did not affect the localization of SPC-1 (Figure S1E, F), suggesting that TWF-2 is not required for spectrin localization. To check if SPC-1 and TWF-2 co-localise, we crossed SPC-1::mKate strain with GFP::TWF-2. Quantitative analysis revealed strong co-localisation between them with a Pearson correlation coefficient of 0.9730 (S1G, H).

### TWF-2 loss of function partially rescues the phenotype of CP knockdown and the two proteins co-localize in some tissues

In other model systems, loss of *twinfilin* leads to structural and functional defects. To investigate whether loss of TWF-2 leads to a phenotype in *C. elegans* we generated a *twf-2* deletion strain using CRISPR/Cas9 (Figure S2A). Surprisingly, this mutant exhibited no discernible phenotype under standard conditions. To evaluate a possible effect on the level or organization of F-actin in the *twf-2* deletion strain, we crossed it with a strain expressing the F-actin-bundling protein plastin/PLST-1 endogenously tagged with GFP. Examination of PLST-1::GFP in embryos did not reveal any difference between the *twf-2* KO and control PLST-1::GFP worms (Figure S2B).

We hypothesized that the lack of phenotype in *twf-2* null worms might be due to its redundancy with other actin dynamics regulators and that its absence might only become apparent when another actin dynamics regulator is depleted. To test this idea, we assembled a list of actin regulators and knocked them down individually by RNAi in the background of *twf-2* null worms (Table S1B). We compared the embryonic lethality of each RNAi in wild-type worms to the lethality in *twf-2* worms. Surprisingly, we did not identify any gene knockdown that resulted in higher embryonic lethality in the *twf-2* null strain as compared to control. In contrast, we found that knockdown of the worm CP CAP-1 had a less severe phenotype in the *twf-2* null background as compared to control. While *cap-1* RNAi led to 47±19% embryonic lethality in wild-type worms it only led to 28±10% embryonic lethality in *twf-2* null worms (Figure 2A).

**Figure 2:**
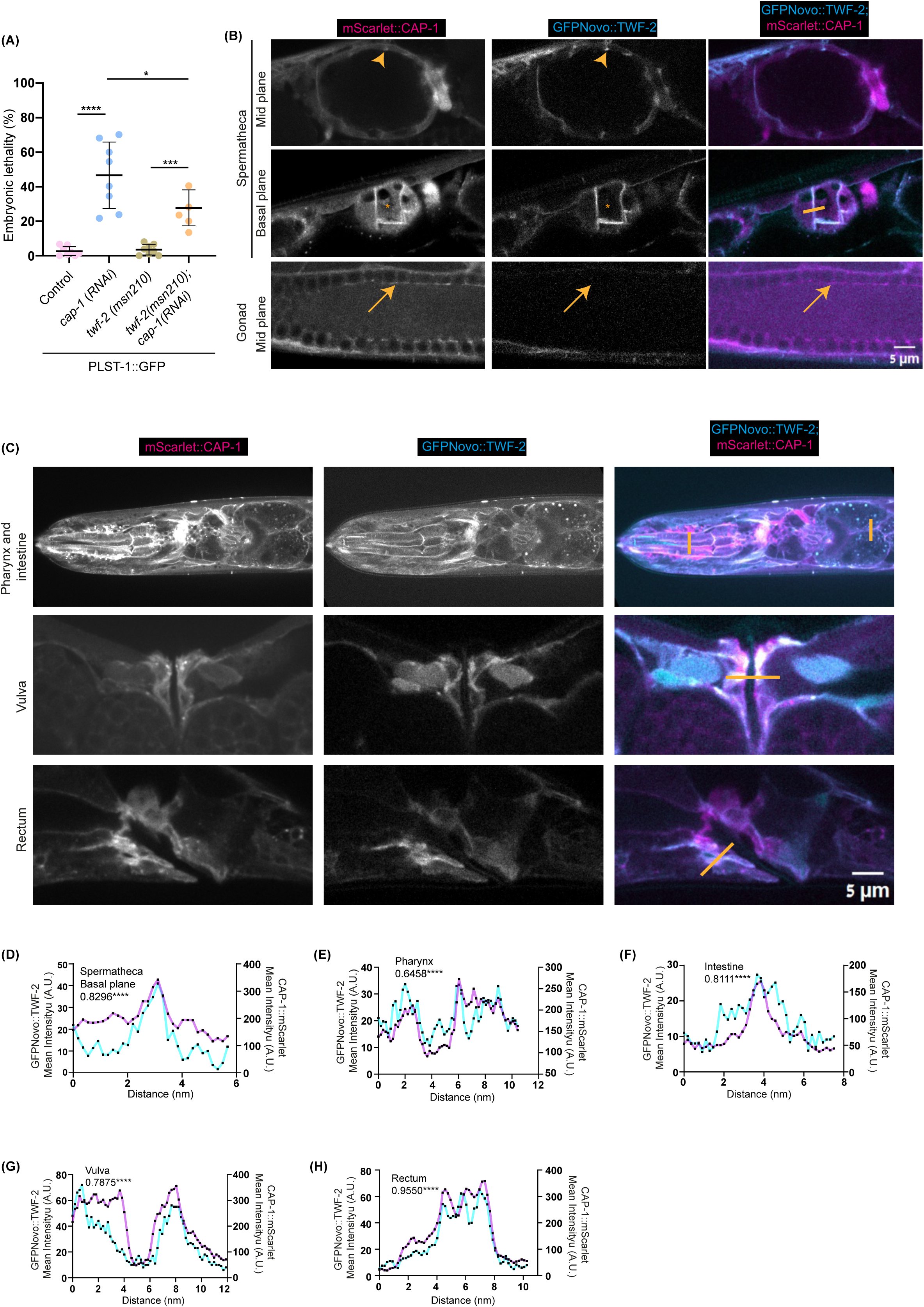
**Genetic interaction between TWF-2 and CAP-1 and their tissue specific colocalization** (A) Embryonic lethality assay. Dot plots show the percentage of embryonic lethality for control (wild type) , *cap-1 RNAi* on wild type, *twf-2(msn210)* mutant control, and *twf-2(msn210); cap-1(RNAi)*. Statistical significance was determined by Ordinary one way ANOVA (ns: not significant, *p < 0.05, **p < 0.01, ***p < 0.001). N=3 biological replicates with ≥298 embryos analysed for each condition. (B) Subcellular co-localization of endogenous GFP::TWF-2 (cyan) and mScarlet::CAP-1 (magenta) in reproductive tissues. Cortical co-localization in spermatheca (arrowheads) contrasts with CAP-1-only regions in spermathecal cytoplasm (asterisks) and germline rachis (arrows). (C) Co-localization of endogenous GFP::TWF-2 (cyan) and mScarlet::CAP-1 (magenta) in other contractile tissues (pharynx, intestine, vulva and rectum) . Orange lines indicate regions used for line profile quantification in (D-H). (D) – (H) Line profile analyses of fluorescence intensity for GFP::TWF-2 (cyan) and mScarlet::CAP-1 (magenta) across indicated regions in (B) and (C). Pearson correlation coefficients are indicated at top left of each graph.

Previous studies in other model systems and *in vitro* experiments have demonstrated that twinfilin and capping proteins often closely associate to regulate actin dynamics (Hakala et al., 2021; Ulrichs & Shekhar, 2025). To investigate whether this association also occurs in *C. elegans*, we crossed the endogenously-tagged mScarlet::CAP-1 strain with the GFP::TWF-2 strain and examined their relative localization. We observed substantial co-localization of CAP-1 and TWF-2 in the spermathecal cortex, pharynx, intestine, vulva and rectum, suggesting a potential functional interaction in these tissues (Figure 2B-H). However, we also observed the localisation of CP in certain tissues where TWF-2 was absent. For example, CAP-1 prominently localized to the germline, while TWF-2 was not detected there (Figure 2B). Moreover, also in tissues where both proteins were expressed they did not always co-localize. For example, CAP-1 was present in the cytoplasm of the spermatheca while TWF-2 was notably absent from this region (Figure 2B). These findings indicate that CAP-1 and TWF-2 are not always required together but instead might participate jointly in some processes and separately in other processes, depending on the context. Furthermore, it appears that TWF-2 is a negative regulator of CAP-1 activity, since its loss alleviated the consequences of *cap-1* RNAi.

### TWF-2 promotes barbed-end depolymerization and is a potent uncapper of CAP-1 from actin filaments

Based on the previous results, we hypothesized that in *C. elegans*, twinfilin is an uncapper of CP. To test this and biochemically characterize the activity of TWF-2, we performed a series of *in vitro* experiments using microfluidics-assisted Total Internal Reflection Fluorescence (mf-TIRF) microscopy (Shekhar, 2017). *C. elegans* TWF-2, CAP-1 and CAP-2 were purified as recombinant proteins from *E. coli*. Building upon previous work demonstrating twinfilin’s actin-depolymerizing activity (Shekhar et al., 2020), we first examined its effects on free barbed ends. Actin filaments with their barbed ends free were elongated from coverslip-anchored spectrin-actin seeds (Figure 3A). The filaments were then aged for 15 minutes to generate ADP-actin filaments. Consistent with previous studies on mammalian twinfilin, we observed that 5 µM TWF-2 significantly decreased the depolymerization rate of aged ADP-actin filaments compared to buffer controls (Figure 3B). Notably, this activity was specific to ADP-actin, as TWF-2 showed an opposite effect by increasing the rate of barbed-end depolymerisation of unaged ADP-P_i_-actin filaments (Figure 3C), suggesting nucleotide-state dependent regulation.

**Figure 3:**
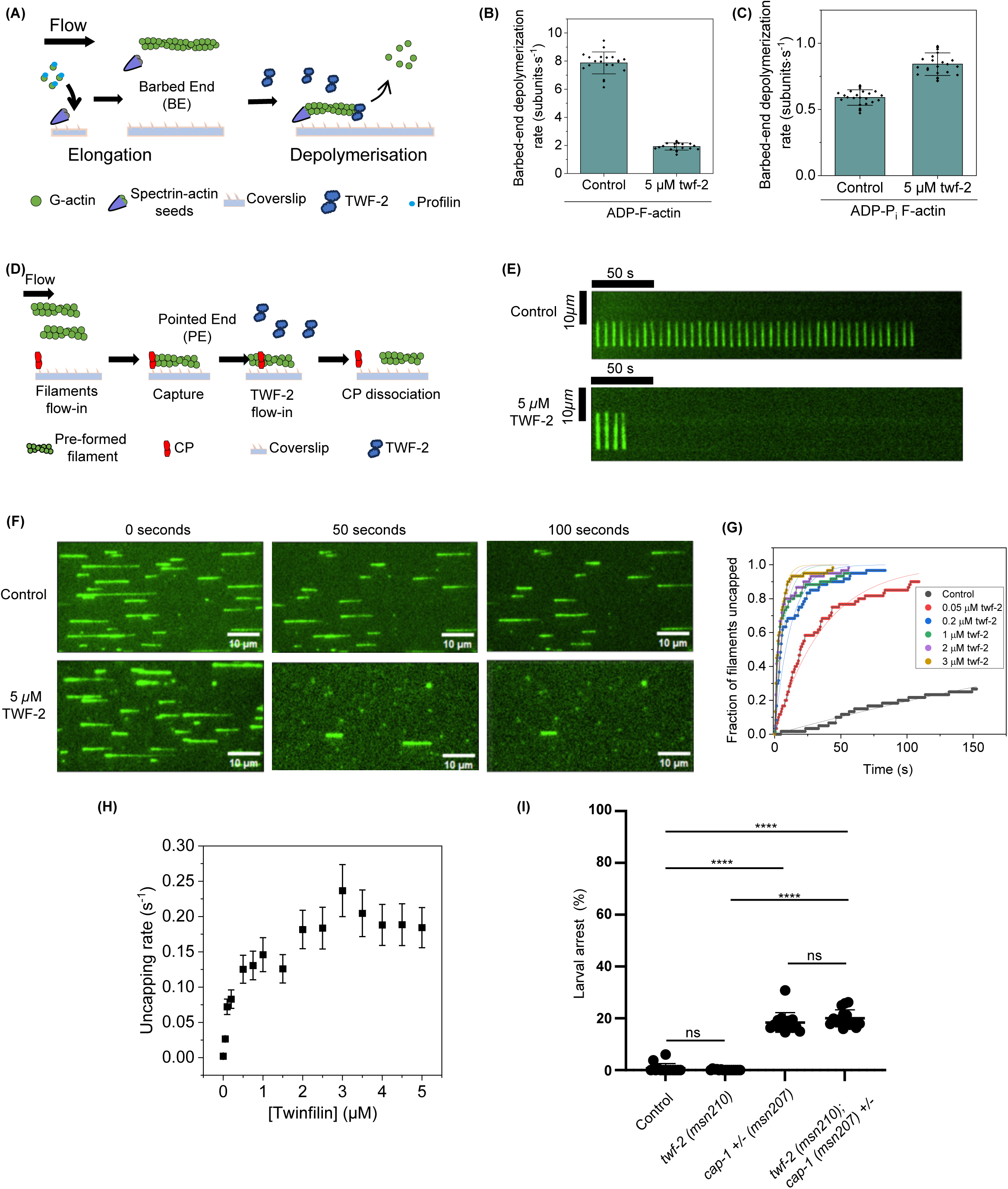
**Effect of TWF-2 on barbed-end depolymerization and uncapping of CP from actin filaments** (A) Schematic representation of the experimental strategy. Actin filaments with free barbed ends were polymerized by exposing coverslip-anchored spectrin-actin seeds to 1 µM G-actin (15% Alexa-488 labeled) and 4 µM profilin. To generate ADP-actin filaments, the filaments were first aged for 15 min first and then exposed to TWF-2 in TIRF buffer. For experiments with ADP-P_i_ filaments, 50 mM excess phosphate was added to all reactions and filaments were not aged. Barbed-end depolymerization was monitored over time. BE, barbed end. (B) Rates (±SD) of barbed-end depolymerization of ADP-actin filaments in the presence of buffer (control) or 5 μM TWF-2. 20 filaments were analysed for each condition. (C) Rates (±SD) of barbed-end depolymerization of ADP-P_i_-actin filaments in the presence of buffer (control) or 5 μM TWF-2. 25 filaments were analysed for each condition. (D) Schematic representation of the experimental strategy. Preformed Alexa-488 labelled ADP-actin filaments were captured by coverslip-anchored biotinylated SNAP-CP. Time-dependent detachment of filaments at a range of TWF-2 concentrations was monitored. BE, barbed end; PE, pointed end. (E) Representative kymographs of a CP-anchored filament uncapping in the presence of buffer (top) or 5 µM TWF-2 (bottom). (F) Representative mf-TIRF time-lapse images showing filament departure due to uncapping. Filaments were exposed to either buffer (top) or 5 µM TWF-2 (bottom). (G) Fraction of filaments uncapped as a function of time in the presence of range of TWF-2 concentrations. Experimental data (symbols) are fitted to a single-exponential function (lines). 59 – 60 filaments were analysed for each condition. (H) CP dissociation rate as a function of TWF-2 concentration, determined from data shown in (G). (I) Larval arrest quantifications. Dot plots indicate percentage larval arrest for control(wild type), *twf-2(msn210) mutant, cap-1(msn207)* mutant and *cap-1(msn207); twf-2(msn210)* double mutant strains (N≥14). Statistical significance was determined by Kruskal Wallis test (ns: not significant, *p < 0.05, **p < 0.01, ***p < 0.001).

We next investigated whether TWF-2 could displace CP from filament barbed ends (Figure 3D). By anchoring biotinylated SNAP tagged *C. elegans* heterodimeric CP to coverslips and capturing preformed Alexa-488-labelled actin filaments, we visualized uncapping events in real-time. Time-lapse imaging revealed that TWF-2 induced rapid filament detachment (Figure 3E, F), with 5 µM TWF-2 uncapping approximately 80% of filaments within 100 seconds (Figure 3F, G). Quantitative analysis demonstrated a concentration-dependent increase in uncapping rates by TWF-2, eventually reaching a saturation level (Figure 3H).

These results provide direct biochemical evidence that *C. elegans* TWF-2 functions both as a barbed-end depolymerizer and as a potent uncapping factor, thereby negating the role of CP. Hence, we hypothesised that the partial rescue of embryonic lethality of *cap-1* RNAi in *twf-2* null worms is because in the absence of the decapper TWF-2, capping activity by the remaining CP is more effective. However, another possibility is that the rescue is indirect, due to the lack of some twinfilin activity that is independent of CP. To distinguish between these two possibilities, we tested whether the absence of twinfilin was able to rescue a complete loss of *cap-1*. For this, we crossed the *twf-2* null strain with a *cap-1* null mutant. The homozygous *cap-1* null is non-viable (100% larval lethal) and is therefore maintained as a balanced heterozygous. After crossing with the *twf-2* null mutant we followed the progeny of the *cap-1/+;twf-2* mothers and found that homozygous *cap-1;twf-2* double mutant progeny, all arrested as larva, similar to the *cap-1* null mutant progeny and did not yield any phenotypic rescue compared to *cap-1* null alone (Figure 3I). Thus, a minimum level of CP is necessary for *twf-2* loss to have an impact on lethality, supporting the notion that the rescue is occurring because TWF-2 works as a decapper of CP in *C. elegans*.

### Loss of *twf-2* alleviates *cap-1* RNAi-associated embryonic lethality by reducing spermathecal contractility

Previously, we characterized the consequences of *cap-1* RNAi in the germline, and showed that it leads to a twofold increase in F-actin and myosin and a hypercontractile phenotype (Ray et al., 2023). However, TWF-2 is not expressed in the germline (Figure 2B). Therefore, the rescue of *cap-1* RNAi-induced embryonic lethality in the *twf-2* null is not likely due to an interaction between them in the germline. On the other hand, we found both capping protein and twinfilin to be highly expressed and co-localized in the spermatheca (Figure 2B), and it is well established that hypercontractility of the spermatheca leads to perturbed embryonic morphology, due to excessive squeezing and occasional pinching, and consequently to embryonic lethality (Tan & Zaidel-Bar, 2015). Therefore, we hypothesized that loss of *twf-2* might rescue the embryonic lethality of *cap-1* RNAi, at least partially, through suppression of spermathecal hypercontractility.

To test this, we performed *cap-1* RNAi in wild-type and *twf-2* worms, and quantified the embryonic morphology. Correlation plots of embryo area and axial ratios revealed significant morphological perturbations in *cap-1* RNAi embryos, consistent with spermathecal dysfunction. Strikingly, *twf-2* deletion partially rescued these defects, reducing the frequency of embryos with extreme morphologies (Figure S3). These findings support the notion that TWF-2 modulates *cap-1* RNAi–associated lethality through its role in spermathecal contractility.

Our data is consistent with the idea that TWF-2, through its uncapping activity, regulates actomyosin contractility in the spermatheca. However, CP is expressed in other tissues and in the embryos, making it difficult to draw conclusions from our results specifically about the spermatheca. In order to circumvent this, we sought conditions of increased contractility specific to the spermatheca. Previously, we showed that knockdown of the RhoGAP SPV-1 leads to ∼40% embryonic lethality due to a hypercontractile spermatheca owing to excess RHO-1 activity (Tan & Zaidel-Bar, 2015). Importantly, *spv-1* is expressed exclusively within the spermatheca. Systemic *spv-1* RNAi in *twf-2* null mutants led to a 63% rescue of embryonic lethality (19.2±8 % in control versus 7±5 % in *twf-2* null) (Figure 4A), suggesting that TWF-2 is required for the increase in spermathecal contractility downstream of RHO-1 activation. Similarly, systemic *twf-2* RNAi in *spv-1* mutants produced comparable results with 54% reduction in embryonic lethality (30.4±13 % in control versus 13.9±11 % in *twf-2 RNAi*) (Figure 4B).

**Figure 4:**
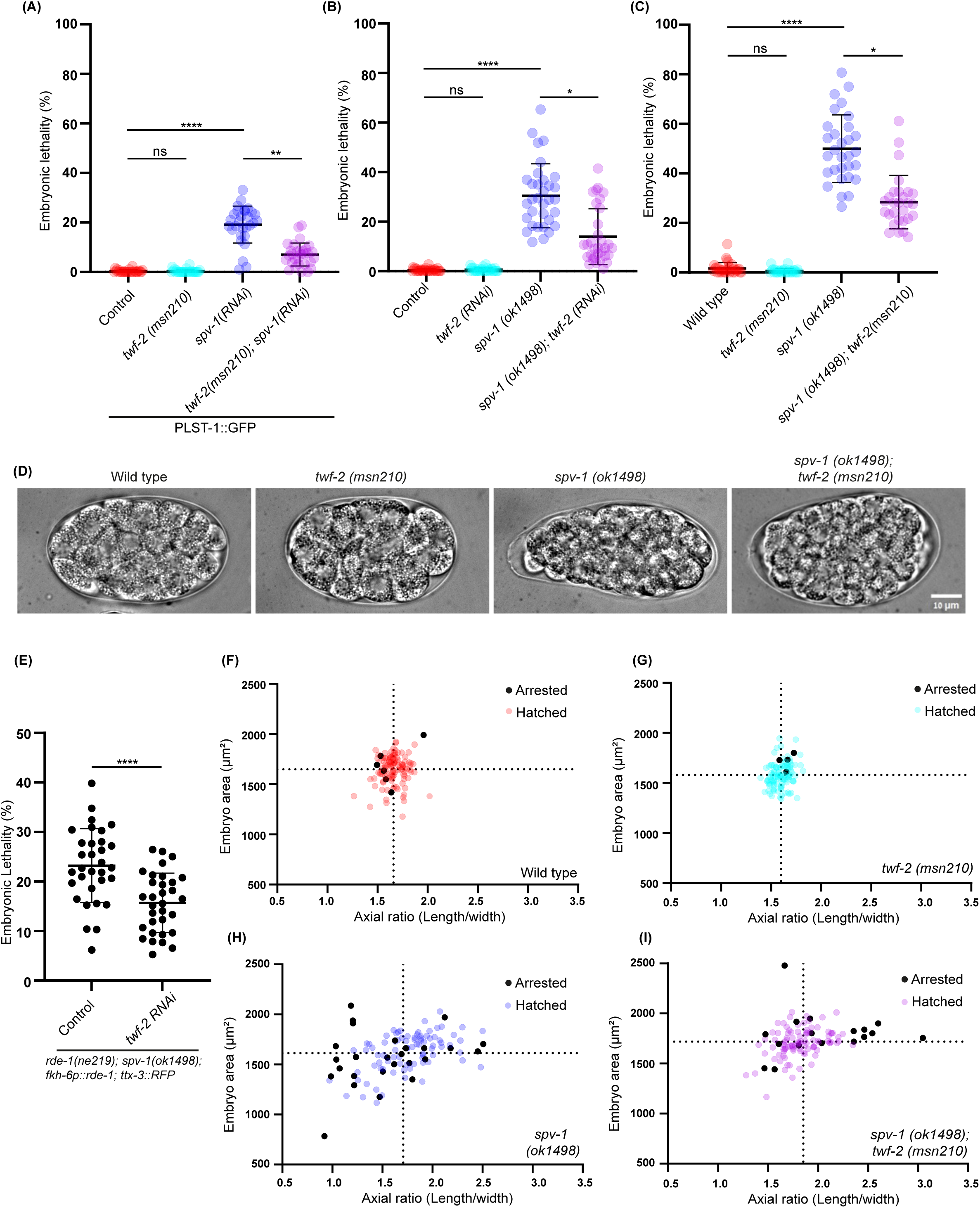
**Genetic interaction between TWF-2 and SPV-1 regulates embryonic viability and morphology** (A) Embryonic lethality in control and *twf-2(msn210)* mutant subjected to systemic *spv-1 RNAi*. Statistical significance was determined by Kruskal Wallis ANOVA test (ns: not significant, *p < 0.05, **p < 0.01, ***p < 0.001). N=3 biological replicates with n≥2700 embryos scored per condition. (B) Embryonic lethality in control and *spv-1(ok1498)* mutant upon systemic *twf-2 RNAi*. Statistical analysis as in (A) (ns: not significant, *p < 0.05, **p < 0.01, ***p < 0.001). N=3 biological replicates with n ≥ 2500 embryos scored per condition. (C) Embryonic lethality in control, *twf-2(msn210)* mutant*, spv-1(ok1498)* mutant *and twf-2(msn210);spv-1(ok1498)* double mutant strains. Statistical analysis as in (A). N=3 biological replicates with ≥2200 embryos scored per condition. (D) Representative DIC images showing embryo morphology for area and axial ratio (length/width) in wild-type, *twf-2 (msn210), spv-1 (ok1498), and twf-2(msn210);spv-1(ok1498)* double mutant strains. (E) Tissue-specific rescue using spermatheca-restricted RNAi. Dot plot shows embryonic lethality in *spv-1(ok1498)* mutant, upon control and *twf-2 RNAi.* Statistical significance was determined by Mann-Whitney test. (****p < 0.001). n≥2500 embryos were scored for each condition. (F) -(H) Quantitative analysis of embryo morphology. Correlation plots show embryo area versus axial ratio in wild-type, *twf-2 (msn210)* strain*, spv-1(ok1498)* and *twf-2(msn210);spv-1(ok1498)* double mutant strains respectively. Black points denote arrested embryos; coloured points denote hatched embryos. Dotted lines indicate the mean values. n≥109 embryos per genotype.

To further validate the genetic interaction between *twf-2* and *spv-1*, we generated a double mutant. This cross confirmed a 43% reduction in embryonic lethality in the double mutant (28.4±11%) compared with the *spv-1* mutant alone (49.96±14%) (Figure 4C). To further confirm that the rescue of embryonic lethality originates from the spermatheca, we performed *twf-2* RNAi in a spermatheca-specific RNAi strain, crossed with the *spv-1* mutant and observed a similar reduction in embryonic lethality phenotype (Figure 4E).

The hypercontractility of *spv-1* mutant spermathecae manifests itself in abnormal embryonic morphology due to abnormal squeezing and pinching of the embryos while they transit through the spermatheca (Tan & Zaidel-Bar, 2015). To investigate whether *twf-2* loss rescued the morphological defects in embryos resulting from *spv-1* loss and the associated hypercontractility of the spermatheca, we analysed embryo shape in all conditions. Remarkably, in the absence of *twf-2*, embryos from *spv-1* mutants exhibited a significant recovery in embryonic morphology, with embryo area and axial ratio parameters returning to near-normal distributions, as visualized in correlation plots (Figure 4 D, F-I).

### *twf-2* loss rescues the increased F-actin levels of the spermatheca in *spv-1 mutants*

To uncover the molecular mechanism underlying the functional interaction between TWF-2 and SPV-1, we examined the actomyosin cytoskeleton in the spermatheca, by performing Phalloidin and phospho-myosin staining on extruded spermathecae from wild-type (N2), *twf-2* mutants, *spv-1* mutants, and *twf-2;spv-1* double mutants. Our analysis revealed that *spv-1* loss caused a significant increase in F-actin and phosphorylated myosin levels in the spermatheca, as expected from its role as a negative regulator of RHO-1 (Figure 5 A-C). Interestingly, *twf-2* loss completely rescued the elevated F-actin levels in the *spv-1* mutants (Figure 5), while it did not affect the elevated levels of phosphorylated myosin (Figure 5). We also checked for the endogenous non muscle myosin-1 (NMY-1) levels in the control, single and double mutant strains and observed no changes with respect to the control (Figure S4). These results strongly suggests that *TWF-2* specifically regulates F-actin levels in the spermatheca, presumably through uncapping of CP, without directly influencing myosin activity.

**Figure 5:**
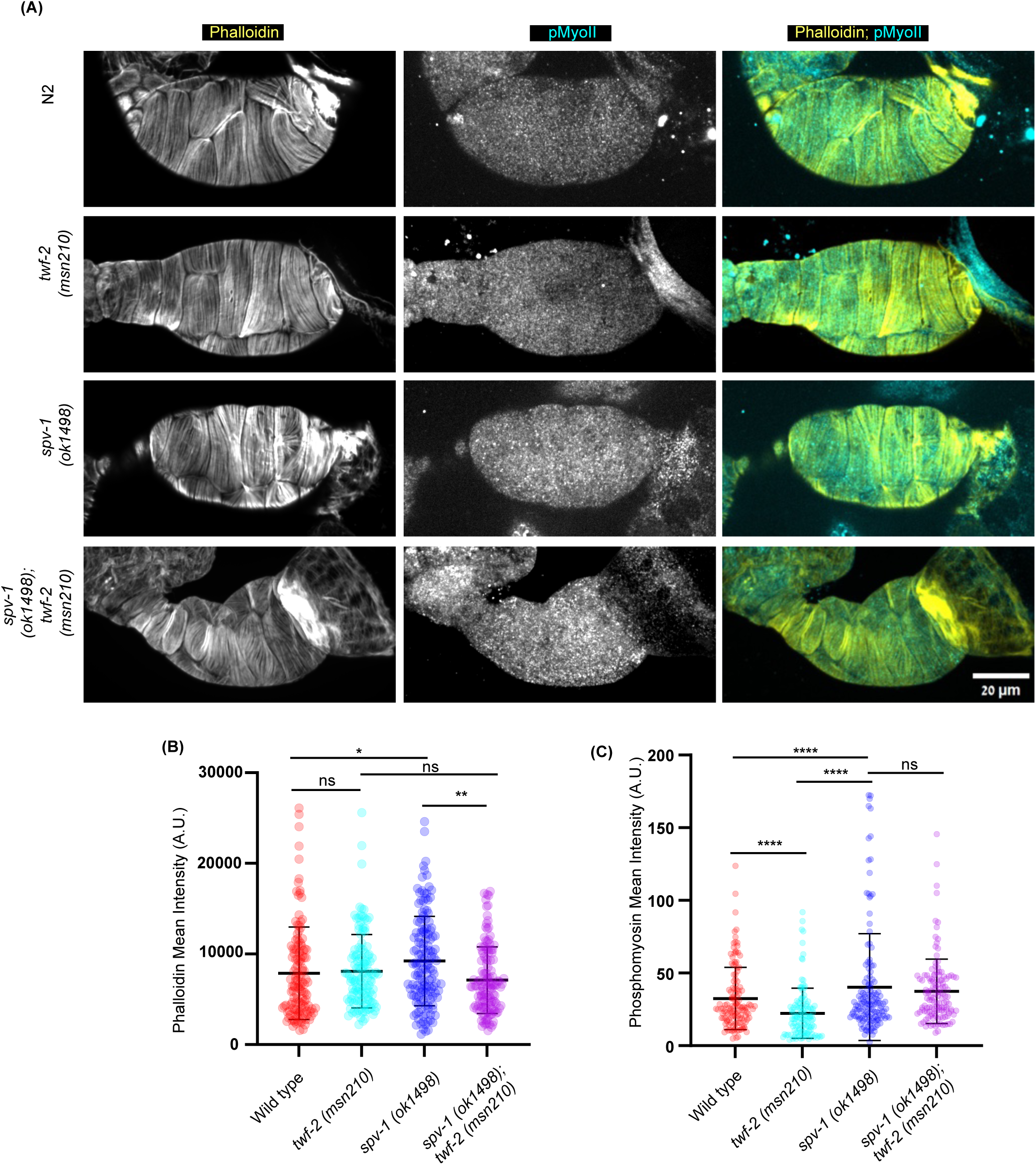
**TWF-2 specifically regulates F-actin levels in the spermatheca** (A) Representative confocal images of extruded spermathecae stained for F-actin (phalloidin, yellow) and phosphorylated myosin (cyan) in wild-type, *twf-2 (msn210), spv-1 (ok1498)*, and *twf-2(msn210);spv-1* double mutant. (B) Quantification of mean fluorescence intensity for F-actin for extruded spermatheca in wild-type, *twf-2 (msn210), spv-1 (ok1498)*, and *twf-2(msn210);spv-1* double mutant. Statistical significance was determined by Kruskal Wallis ANOVA test (ns: not significant, *p < 0.05, **p < 0.01, ***p < 0.001). n≥123 spermathecae per genotype from 3 biological replicates were quantified. (C) Quantification of mean fluorescence intensity for Phosphomyosin for extruded spermatheca in wild-type, *twf-2 (msn210), spv-1 (ok1498)*, and *twf-2(msn210);spv-1* double mutant. Statistical significance was determined as in (B). n≥123 spermathecae per genotype from 3 biological replicates were quantified.

## Discussion

In this study, we provide the first comprehensive characterization of *C. elegans* twinfilin-2 (TWF-2), the sole twinfilin ortholog in this organism, revealing its broad expression and its role as a modulator of spermathecal contractility through CP uncapping. We demonstrate that TWF-2 exhibits cortical localization, which in the spermatheca is dependent on α-spectrin (SPC-1) and β-spectrin (UNC-70). Interestingly, while we identified spectrin in a pulldown of twinfilin, a previous IP-MS study (Jia et al., 2020) identified TWF-2 (annotated as F38E9.5) as a potential SPC-1 interactor, further supporting their association. Whether they interact directly or indirectly is a question that requires further experiments to answer. The dependence of TWF-2 cortical localization on spectin, only in the spermatheca, highlights specialized mechanisms for spatial control of actin dynamics in distinct tissues.

While TWF-2 and CP functionally interact in regulating spermatheca contractility, as manifested in changes in embryo morphology, their localization patterns do not always align. Their partial dissociation implies context-specific roles. A known TWF-2-independent function of CP is in the dynein-dynactin complex. TWF-2 might have CP-independent functions, yet to be discovered. These observations underscore the multifunctional nature of both proteins, with their interactions being precisely regulated in a tissue- and process-dependent manner.

Our *in vitro* results revealed *C. elegans* TWF-2’s conserved ability as an uncapper, with its barbed end depolymerisation activity, dependent on the F-actin nucleotide state, being similar to mouse twinfilin (Shekhar et al., 2020). The concentration-dependent uncapping kinetics found *in vitro* suggest that the differential expression levels of TWF-2 we observed in specific tissues are significant toward TWF-2’s uncapping activity *in vivo*. Notably, whereas mouse twinfilin enhances uncapping by only sixfold (Hakala et al., 2021; Reddy et al., 2025), *C. elegans* twinfilin accelerates it by over 90-fold, suggesting a species-specific evolutionary adaptation for rapid uncapping. The uncapping activity *in vitro* also provides a mechanistic basis for our genetic observations *in vivo*, where the partial rescue of *cap-1*(RNAi) embryonic lethality by loss of *twf-2*, with respect to spermatheca hypercontractility, demonstrates that TWF-2, in this context, antagonises CP function. Absence of rescue in double mutant of *cap-1;twf-2* confirms the direct relationship between the two, with a requirement of minimal CP activity for *twf-2* null to rescue loss of CP.

The physiological relevance of TWF-2-mediated actin regulation specifically in the spermatheca is highlighted by its genetic interaction with the RhoGAP SPV-1, which is only expressed in the spermatheca. We found that *twf-2* loss mitigates the embryonic lethality and morphology defects of *spv-1* loss while completely rescuing the increased F-actin levels found in this mutant. Importantly, *twf-2* loss did not rescue the increase in non-muscle myosin II expression and phosphorylation levels found in the *spv-1* mutant, reflecting a role for TWF-2 strictly related to actin and not myosin. This is the opposite of what has been reported for mouse cochlear mechanosensory stereocilia (Rzadzinska et al., 2009). This uncoupling of actin and myosin regulation suggests TWF-2 may serve as a specialised modulator of actin assembly in response to mechanical cues, such as ovulation events.

Our findings bridge the gap between *in vitro* biochemistry and tissue-level function, showing how TWF-2’s molecular activities (uncapping, nucleotide-state-sensitive depolymerization) translate to physiological control of contractility. The spectrin-TWF-2-CP axis we describe is potentially relevant to other contractile systems, primarily at the cell cortex. The *C. elegans* spermatheca, with its genetic accessibility and mechanical readouts, provides an ideal system to address questions related to actin binding proteins in contractility. More broadly, our work establishes TWF-2 as a key node linking spectrin-based cortical organization, CP-mediated capping, and RHO-1-dependent contractility, offering new insights into the coordination of structural and dynamic elements in actomyosin networks.

## Materials and Methods

### Worm Strain Maintenance

All worm strains were maintained at 15° C, as per standard worm protocols (Brenner, 1974). All the experiments were performed at 20°C. All strains used in this study have been listed in Table S2.

### CRISPR-Cas9-Mediated Generation of Reporter and Knockout Strains

The TWF-2 endogenous reporter strain and *twf-2* null mutant were generated using CRISPR-Cas9-mediated genome editing (Paix et al., 2017). To create an endogenous reporter strain, *twf-2* was tagged at the N-terminus with GFPNovo. A single guide RNA (sgRNA) (aggagaagaagctactactA) was designed to insert the GFPNovo fluorophore, along with a linker sequence (TCTGGTGGTAGTGGCGGTACC), downstream of the 5′ UTR and in frame with the first exon. The repair template, consisting of GFPNovo and the linker sequence, was PCR-amplified from a plasmid and included 35 bp homology arms flanking the 5′ UTR and the first exon. The template was purified using the Macherey-Nagel kit and eluted in water.

The injection mix contained Cas9 protein (0.8 µg/µl), *twf-2* sgRNA (0.0625 nmol/µl), GFPNovo repair template (4000 ng), *dpy-10* sgRNA (0.02 µg/µl), *dpy-10* ssODN (0.05 µg/µl), KCl (25 mM), tracrRNA (0.1 µg/µl), and water in a total volume of 20 µl. The mix was microinjected into the gonads of young adult *N2* hermaphrodites.

F1 progeny with the dumpy (*dpy-10*) phenotype were isolated onto individual NGM plates, allowed to lay embryos, and then genotyped via single-worm lysis followed by PCR. Genotyping was performed using primers (ccgaacatttggatgggaagg and agtcaacacccacttaacccc). The insertion of GFPNovo was confirmed through DNA sequencing.

For generation of null mutant, two sgRNAs were designed to delete a region of 2840 bp region of the *twf-2* gene. The first sgRNA (CTTCGAAATGCACTCAACTT) targeted exon 2 while the second sgRNA (aaattaaatattgcagATGG) targeted exon 8. A single-stranded oligodeoxynucleotide (ssODN) repair template (tttgcagCTAACGCGGCACTTCGAAATGCACTCAATGGAGGTAGATGCTCGCGACGATCTTTCGGAGAAA) was synthesized with 35bp homology arms flanking the deletion site.

The injection mix contained Cas9 protein (0.8ug/ul), *twf-2* sgRNAs (0.0625 nmol/ul), *twf-2* ssODN (0.225ug/ul), *dpy-10* sgRNA (0.02ug/ul), *dpy-10* ssODN (0.05ug/ul), KCl (25mM), tracrRNA(0.1ug/ul), and water in a total volume of 20ul. This mix was microinjected into the gonads of young adult N2 hermaphrodites.

F1 progeny exhibiting the dumpy (*dpy-10*) phenotype were isolated onto individual NGM plates and allowed to lay embryos. These F1s animals were lysed and genotyped by PCR to screen for the deletion. Specific primers (gtcccagacacacttctcttcc and tcgggtcgacgtttcagtatg) were used to identify mutants, while primers (CTCGTTGCCATCATCTGGAAG and tcgggtcgacgtttcagtatg) were used to confirm the wild-type genotype. The targeted deletion was further validated by DNA sequencing of the progeny.

### RNAi knockdown

Systemic and spermatheca-specific RNAi-mediated gene knockdown were performed by feeding *Escherichia coli* HT115 bacteria expressing double stranded RNA (dsRNA) targeting the gene of interest. RNAi clones *spv-1* (T444T) and *twf-2* (L4440) clones were used, with their corresponding empty vector clones (T444T and L4440) as negative controls. RNAi plates with prepared with nematode growth medium (NMG) containing 1mM IPTG and 100ug/ml ampicillin.

Primary cultures of HT115 clones were grown in 5 ml Luria-bertani (LB) broth containing 100ug/ml ampicillin, overnight at 37°C. Secondary cultures were inoculated by adding 200 µl of the primary culture into 20 ml LB containing 100 µg/ml ampicillin and incubated for 7–8 hours at 37°C until the culture reached an optical density (OD_600_) of 1. The bacterial culture was centrifuged, and the pellet was resuspended in 2ml M9 buffer. RNAi plates were seeded with 150 µl of the bacterial suspension and allowed to dry overnight at room temperature.

Adult hermaphrodites of the required genotype were bleached, and embryos were dropped onto the RNAi-seeded plates, after adding 100ul 1M IPTG on the seeded bacteria to induce dsRNA expression. All RNAi experiments in this study were performed at 20°C. Phenotypic analysis was performed on either the same generation or their progeny.

### Embryonic lethality, larval arrest and embryonic shape assays

Embryonic lethality assays were performed using day1 adult hermaphrodites. Ten worms were placed on each RNAi-seeded plates or OP50 seeded plates for 3-4 hours, yielding an average of approximately 100 embryos per plate. After the egg-laying period, the adults were removed, and the total number of embryos were counted. The plates were incubated at 20°C for 24 hours. Following incubation, the embryos were scored for viability by counting the number of hatched and unhatched embryos. Larval arrest was scored after an incubation of 48 hours.

Embryo shape and area were analysed by dissecting day 1 adult hermaphrodites. The embryos were mounted on 3% agarose pads prepared on glass slides, sealed with wax after the addition of M9 buffer. Each embryo was numbered and imaged for shape analysis.

Following imaging, the slides were incubated overnight at 20°C. Embryo viability was assessed 12–16 hours post-incubation by scoring hatching.

### Phalloidin staining

Worms were collected in M9 and extruded after addition of 100mM Levamisole, within 5 minutes. Fixation was performed by adding 4% formaldehyde in PBS, for 15 minutes. The formaldehyde was removed was centrifugation at 1500 rcf for 30 seconds and cold acetone (-20°C) was added for 5 minutes.

The samples were washed thrice with PBST (1X PBS containing 0.5% Triton X-100). Phalloidin conjugated to TRITC was added to a final dilution of 1:250 in PBST and incubated at room temperature for 2 hours. Post incubation, the samples were washed thrice with PBST, and DAPI was added to a dilution of 1:1000 in PBST, followed by a 30-minute incubation at room temperature.

The samples were washed again with PBST and Vectashield mounting media was added to the samples which were either stored at 4°C for up to a week or immediately mounted for imaging.

### Immunofluorescence

Worm gonads were extruded in M9 buffer containing 100mM Levamisole within 5 minutes. The dissected gonads were fixed with 4% formaldehyde in PBS for 15 minutes, followed by three washes in PBS. Permeabilization was performed using 0.25% Tween 20 in PBS for 10 minutes, followed by three washes thrice with PBS again.

The samples were incubated in a blocking solution comprising 1% bovine serum albumin(BSA), 0.1% Tween 20 and 30mM glycine in PBS for 1 hour at room temperature. Primary antibody incubation was carried out overnight at 4°C using anti-phospho-MLC (Ser19) (Cell Signalling Technology, #3671) at a 1:400 dilution in the blocking solution.

Gonads were washed thrice with PBS and incubated with blocking buffer containing 1:500 anti-rabbit secondary antibody conjugated with Alexa 488 (Invitrogen, A21244), 1:250 phalloidin-TRITC (Sigma-Aldrich, P1951) and 1:1000 DAPI (Sigma-Aldrich), at room temperature for 90 minutes. Following three washes with PBS, Vectashield mounting media (Vector Laboratories, H-1000) was added. The gonads were stored at 4°C for up to a week or directly mounted for imaging.

### Image acquisition

Imaging was performed using a Nikon Ti-2 Eclipse inverted microscope equipped with a Yokogawa spinning-disk confocal system (CSU-W1) and Plan-Apochromat oil-immersion objectives (60×, 1.4 NA or 100×, 1.45 NA). Samples were illuminated with 405 nm, 488 nm, and 561 nm lasers (Gataca Systems) and captured using a Prime 95B sCMOS camera (Photometrics). MetaMorph software (Molecular devices) was used for acquisition control. All imaging was performed at 20°C.

For fluorescence and differential interference contrast (DIC) microscopy, worms were mounted on 3% agarose pads and immobilized using Levamisole. Embryos were mounted in M9 buffer and imaged using 100X 1.45NA oil-immersion objective.

### Image analysis and Statistics

All image analyses were performed using ImageJ (NIH)(Schindelin et al., 2012). Regions of interest (ROIs) were manually selected for subsequent quantification. For co-localization analysis (Figure 1), a 5-pixel-wide line was drawn along the region of interest for both channels. Fluorescence intensity was normalized and plotted on a graph.

Embryo shape and area were quantified by manually tracing the eggshell using the segmented line tool in ImageJ. Embryo length and width were measured by drawing straight-line segments along the longest and shortest axes, respectively.

Phalloidin quantifications were performed by drawing a 20-pixel-wide line at a specific z-stack to measure the mean intensity of actin fibres in a single cell. A maximum of two cells were analysed per individual spermatheca. Phosphomyosin mean intensity was measured in the same region of interest, using the same 20-pixel-wide line as for the Phalloidin measurements. For both stainings, background intensity was subtracted for each image to ensure accurate quantification. Statistical analysis was performed using the Prism software (GraphPad).

### Immunoprecipitation and Mass Spectrometry

Synchronized GFPNovo::TWF-2 knock-in strains and N2 L1 larvae were cultured on 10 × 60-mm NGM plates at 20°C. After 50–52 hours, mixed populations were collected and washed three times with M9 buffer (42.33 mM Na_2_HPO_4_, 22.06 mM KH_2_PO_4_, 85.56 mM NaCl, 1mM MgSO_4_). Worms were fixed with 0.5% paraformaldehyde (PFA) in M9 buffer for 20 minutes at 20°C. The fixative was removed, and worms were incubated with 50 mM Tris (pH 8.0) in M9 buffer for 5 minutes at 20°C. After two washes with M9, the worm pellet was stored at −80°C.

Frozen samples were thawed on ice and washed with cold 1× PBS. Worms were resuspended in ∼500 µL of lysis buffer (50 mM Tris-HCl pH 7.4, 150 mM NaCl, 1 mM EDTA, 1% Triton X-100) containing a protease inhibitor tablet. Sonication was performed on ice until worm bodies were no longer visible in the buffer. The lysate was centrifuged at 15,000 rpm for 20 minutes at 4°C, and the supernatant was transferred to new tubes. A 30 µL aliquot was set aside for Bradford analysis and Western blotting, while the remaining lysate was stored at −80°C.

Immunoprecipitation was carried out using G-Sepharose beads. For pre-clearance, 15 µL of protein G beads were added to the lysate and incubated at 4°C for 2 hours. After centrifugation at 12,000 rcf for 15 minutes at 4°C, the supernatant was transferred to fresh tubes and incubated overnight with 8 µg of anti-GFP mouse antibody (Roche 11814460001) at 4°C. Following antibody binding, 15 µL of beads were added, and the samples were incubated at 4°C for 3 hours. Beads were collected by centrifugation at 12,000 rcf for 30 seconds at 4°C and washed three times with 1× wash buffer (0.05 M Tris-HCl pH 7.4, 0.15 M NaCl). A final wash was performed with PBS containing 1% Triton X-100.

The samples were boiled in 1× Laemmli sample buffer at 95°C for 10 minutes. After centrifugation at 15,000 rpm, the supernatant was transferred to clean tubes. Ten percent of the sample was used for SDS-PAGE Western blotting as a quality check, and the remaining sample was subjected to Western blot followed by trypsin digestion and analysis by LC-MS/MS on Q-Exactive HF (Thermo). Mass spectrometry was performed at the Smoler Protein Research Center at the Technion Institute of Technology.

### Protein purification for in-vitro experiments

Proteins were purified as per the established protocol (Shekhar et al., 2020). Briefly, rabbit skeletal muscle actin was purified from acetone powder (PelFreez). Lyophilized powder was sheared, resuspended in G-buffer (5 mM Tris-HCl pH 7.5, 0.5 mM DTT, 0.2 mM ATP, 0.1 mM CaCl₂), cleared by centrifugation (50,000 × g, 20 min), and filtered. Actin was polymerized overnight at 4°C by adding 2 mM MgCl₂ and 50 mM NaCl, and then 0.6 M NaCl was added. F-actin was centrifuged, homogenized, dialyzed against G-buffer (48 h), gel-filtered (Sephacryl S-200), and stored at 4°C.

For labelling, G-actin was polymerized in modified F-buffer (20 mM PIPES pH 6.9, 100 mM KCl, 0.2 mM CaCl₂, 0.2 mM ATP), incubated with Alexa-488 NHS ester (5:1 molar excess, 2 h, RT), pelleted (450,000 × g, 40 min), depolymerized in G-buffer, and repolymerized (100 mM KCl, 1 mM MgCl₂). After a final ultracentrifugation, labelled actin was dialyzed, cleared (450,000 × g, 40 min), and assessed for concentration and labelling efficiency.

*C. elegans* TWF-2 (6×His-tagged) was expressed in *E. coli* BL21 (pRare) induced with 1 mM IPTG (18°C, overnight). Cells were lysed in 50 mM NaPO₄ (pH 8), 300 mM NaCl, 20 mM imidazole, 1 mM DTT, and protease inhibitors, sonicated, and cleared (120,000 × g, 45 min). The lysate was incubated with Ni-NTA beads (2 h, 4°C), washed, and eluted with 250 mM imidazole. Purified protein was further resolved by size-exclusion chromatography (Superdex 75 Increase, 20 mM HEPES pH 7.5, 50 mM KCl, 1 mM DTT), concentrated, flash-frozen, and stored at −80°C.

Human profilin-1 was expressed in *E. coli* BL21 (pRare), induced with 1mM IPTG. Cells were lysed in 50 mM Tris-HCl (pH 8), 1 mM DTT, and protease inhibitors, sonicated, and centrifuged (120,000 × g, 45 min). The supernatant was applied to poly-L-proline beads, washed (10 mM Tris pH 8, 150 mM NaCl, 1 mM DTT), and eluted with 8 M urea. The protein was dialyzed (2 mM Tris pH 8, 0.2 mM EGTA, 1 mM DTT), ultracentrifuged (450,000 × g, 45 min), concentrated, flash-frozen, and stored at −80°C.

*C. elegans* SNAP-tagged capping protein was expressed in E. coli BL21 DE3 (1 mM IPTG, 18°C, overnight). Cells were lysed in 20 mM NaPO₄ (pH 7.8), 300 mM NaCl, 15 mM imidazole, 1 mM DTT, and protease inhibitors, sonicated, and cleared (150,000 × g, 30 min). The lysate was purified via HisTrap chromatography, eluted with 250 mM imidazole, and labeled with benzylguanine-biotin. Free biotin was removed by size-exclusion chromatography (Superose 6, 20 mM HEPES pH 7.5, 150 mM KCl, 0.5 mM DTT). Purified protein was aliquoted, flash-frozen, and stored at −80°C.

### Microfluidics-assisted TIRF (mf-TIRF) Microscopy

mf-TIRF microscopy was performed as per the protocol established (Shekhar et al., 2020). Briefly, actin filaments were assembled in mf-TIRF flow cells (Shekhar, 2017), Coverslips were cleaned by sonication in Micro90 detergent and 1 M KOH, 1 M HCl, and ethanol, then coated with mPEG-silane (2 mg/mL) and biotin-PEG-silane (2 µg/mL) in acidic 80% ethanol (pH 2.0) and incubated overnight at 70°C. A PDMS flow chamber (40 µm height, 3 inlets, 1 outlet) was clamped onto the coated coverslip and connected to a microfluidic flow-control system (Fluigent) and rinsed with TIRF buffer. Chambers were incubated with 1% BSA and 10 µg/mL streptavidin in HEPES/KCl buffer.

For uncapping experiments, biotinylated capping protein (CP) was anchored to the coverslip surface, and preformed Alexa-488-labeled actin filaments (15% labeled) were introduced. Filament survival was monitored in TIRF buffer (10 mM imidazole pH 7.4, 50 mM KCl, 1 mM MgCl₂, 1 mM EGTA, 0.2 mM ATP, 10 mM DTT) with and without addition of twinfilin. The cumulative distribution functions were fit to an exponential decay function to determine the barbed-end dissociation rate of CP.

For depolymerization assays, filaments were grown from spectrin-actin seeds using 1 µM G-actin and 4 µM profilin in TIRF buffer, aged to ADP-actin (15 min in 0.1 µM G-actin in TIRF buffer) and then exposed to TIRF buffer with or without twinfilin. Depolymerization rates were measured from kymographs (Fiji). For ADP-Pi-actin experiments, filaments were maintained in phosphate-supplemented TIRF buffer (34.8 mM K₂HPO₄, 15.2 mM KH₂PO₄).

Images were acquired using A Nikon Ti2000 microscope (60× 1.49 NA TIRF objective, Perfect Focus System) with 488/561/640 nm lasers and an EMCCD camera (Andor Ixon 888) was used. Images were acquired with Nikon Elements, drift-corrected (Fiji), and analyzed via kymographs. Rates were averaged from at least three replicates.

## Acknowledgments

Some strains were provided by the CGC, which is funded by NIH Office of Research Infrastructure Programs (P40 OD010440). This work was supported by grant No. 767/20 from the Israeli Science Foundation to RZB and NIH NIGMS grant R35GM143050 to SS.

## Supplementary

**Table S1.** List of RNAi candidates.

|  | Gene names | Source of shortlisting | Readout for screening |
| --- | --- | --- | --- |
| A | *unc-27, hum-1, ule-3, arx-6, unc-52, rab-1, let-60, clik-1, ifb-2, spc-1, unc-70, unc-87, ifc-2, nmy-1, act-5, erm-1, unc-60, mlc-4* | IP-MS | GFPnovo::TWF-2 localisation<br>Embryonic lethality with wild type and *twf-2(msn210) mutant* |
| B | *cyk-1, aipl-1, unc-78, cas-1, cas-2, cap-1, mel-11, gsnl-1, unc-34, cdap-2, unc-60, tth-1, mtpn-1, crml-1, cdap-2* | Actin-interacting proteins | Embryonic lethality with wild type and *twf-2(msn210) mutant* |

**Table S2.**
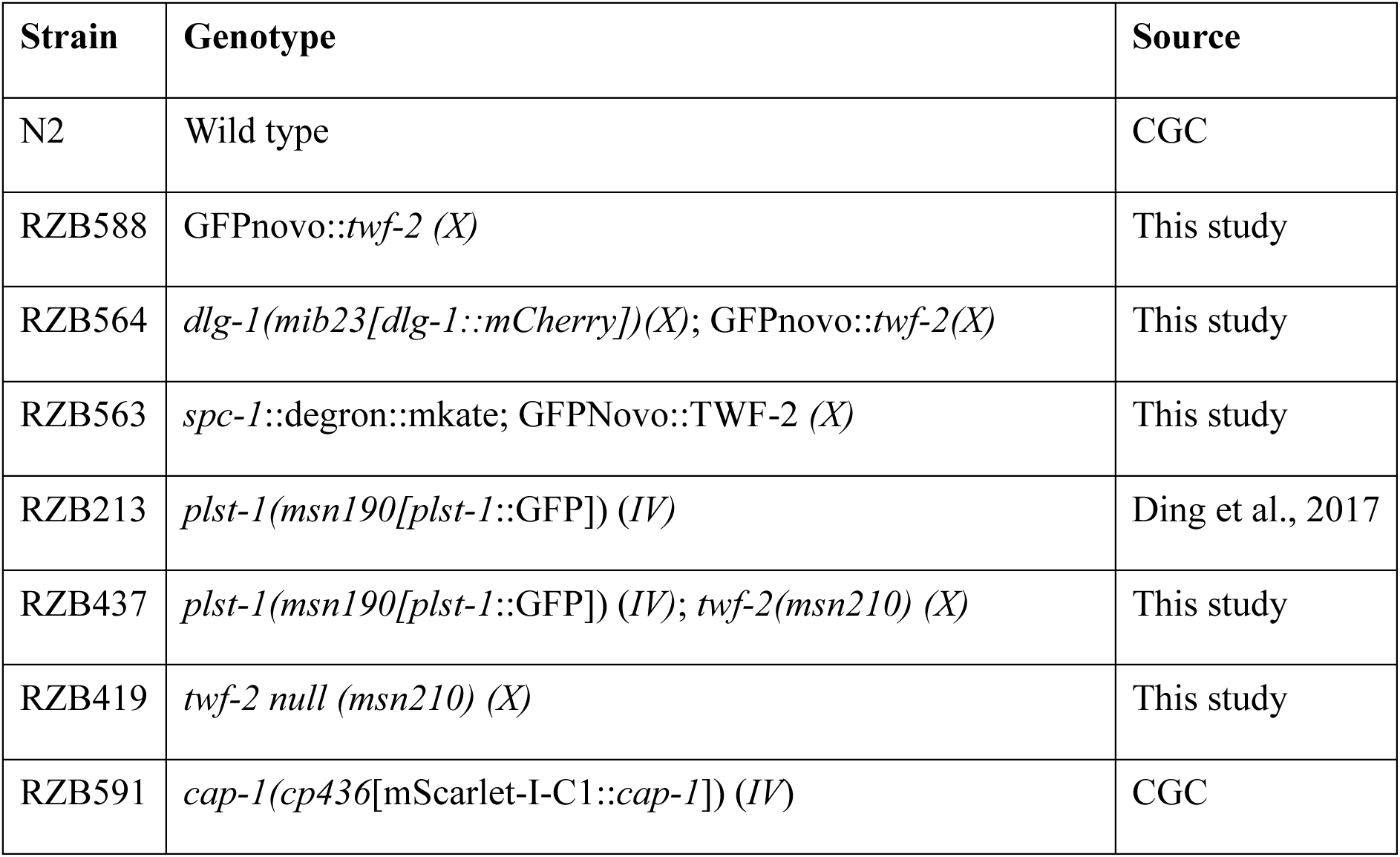
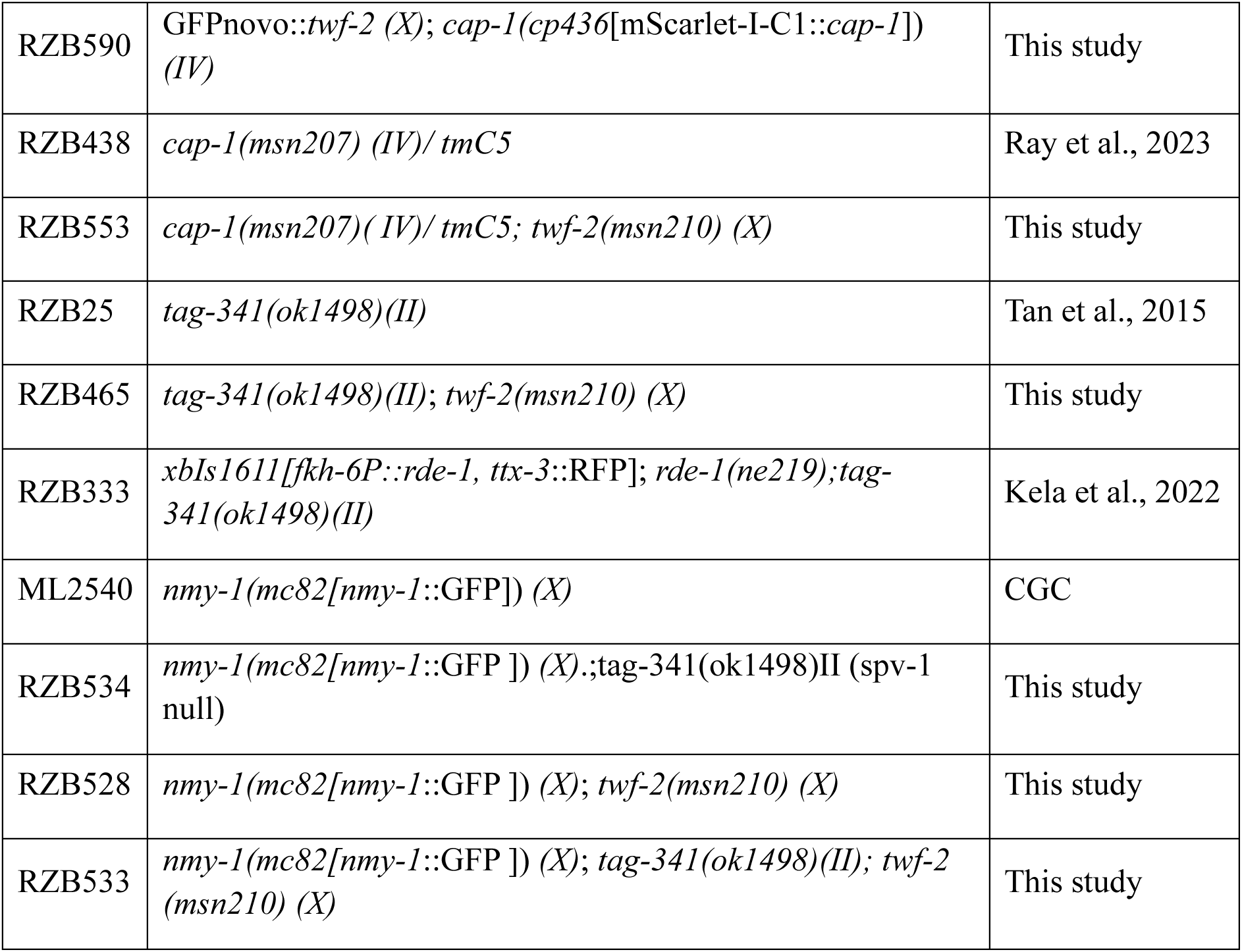
List of all strains used.

---

**Supplementary Figure 1:**
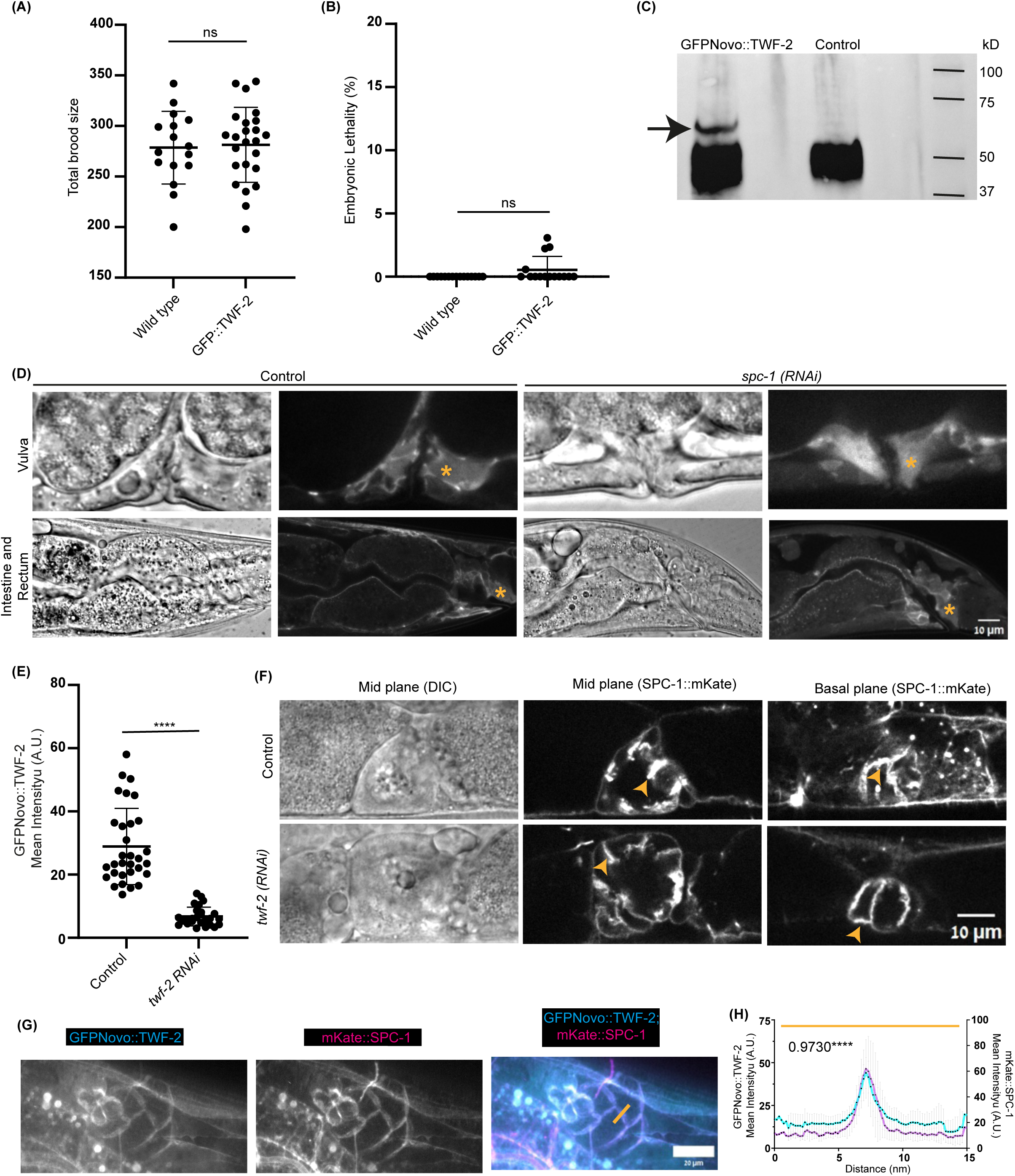
**Characterisation of GFP::TWF-2 strain** (A) Brood size quantification for wild-type (N2) versus GFP::TWF-2 knock-in strains. Each point represents one hermaphrodite (N≥15 worms for each genotype). Statistical significance was determined by Mann-Whitney test. Horizontal bars indicate mean ± SD. (B) Embryonic lethality rates in wild-type versus GFP::TWF-2 animals. Statistical significance was determined by Mann-Whitney test. Horizontal bars indicate mean ± SD (n ≥1000 embryos were scored for each strain) (C) Western blot of GFP::TWF-2 and wild type control strain. From left: GFP::TWF-2 strain; N2 control (untagged). Arrow indicates full-length GFP::TWF-2 fusion protein. (D) Tissue-specific effects of systemic *spc-1(RNAi)* on GFP::TWF-2 localization. Single Z-projections show vulva, intestine, and rectum in control versus *spc-1(RNAi)* animals. Cortical signal is maintained (Asterisks) in all tissues except spermatheca (Fig. 1C). (E) RNAi efficiency quantification. Bar graph shows GFP::TWF-2 fluorescence intensity in spermathecae after control versus *twf-2(RNAi)* (****p < 0.0001, Mann-Whitney test). n≥27 animals were quantified for each condition. (F) SPC-1::mKate localization is unaffected by *twf-2(RNAi)*. Representative images show spermathecal SPC-1 distribution (arrowheads) in control and *twf-2(RNAi)* animals. (G) Co-localization sum projection of Z slices demonstrate spatial overlap between GFP::TWF-2 (cyan) and SPC-1::mKate(magenta) in spermatheca. (H) Co-localization quantification for GFP::TWF-2 and SPC-1::mKate. Line profile plot shows Pearson correlation coefficient of 0.9730.

**Supplementary Figure 2.**
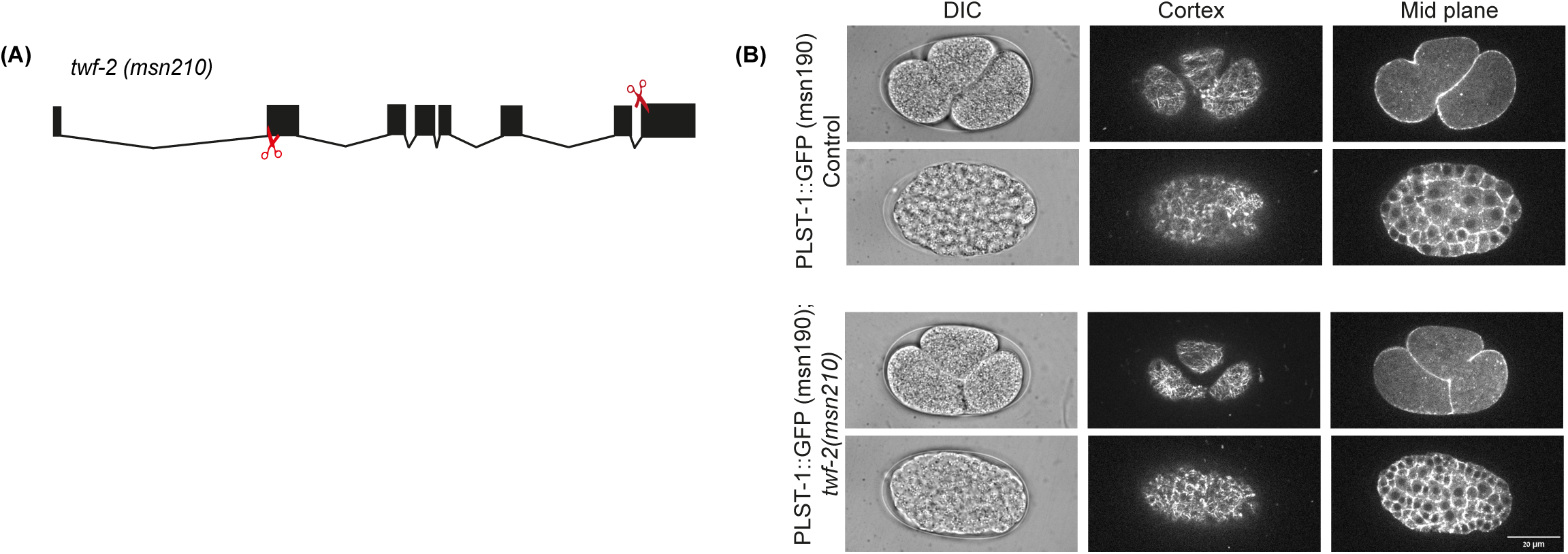
**Generation and validation of *twf-2* null mutant** (A) CRISPR-Cas9 strategy for *twf-2(msn210)* deletion allele. Schematic shows genomic organization of twf-2 locus with exons (black boxes) and introns (connecting lines). Red scissors indicate sgRNA target sites flanking the deletion region. (B) Actin organization in *twf-2(msn210) null* mutants. Representative confocal images of PLST-1::GFP-labelled actin in 3- and multi-celled stage wild-type and *twf-2(msn210)* mutant embryos. Single Z-slices show comparable actin cytoskeleton organization between genotypes.

**Supplementary Figure 3.**
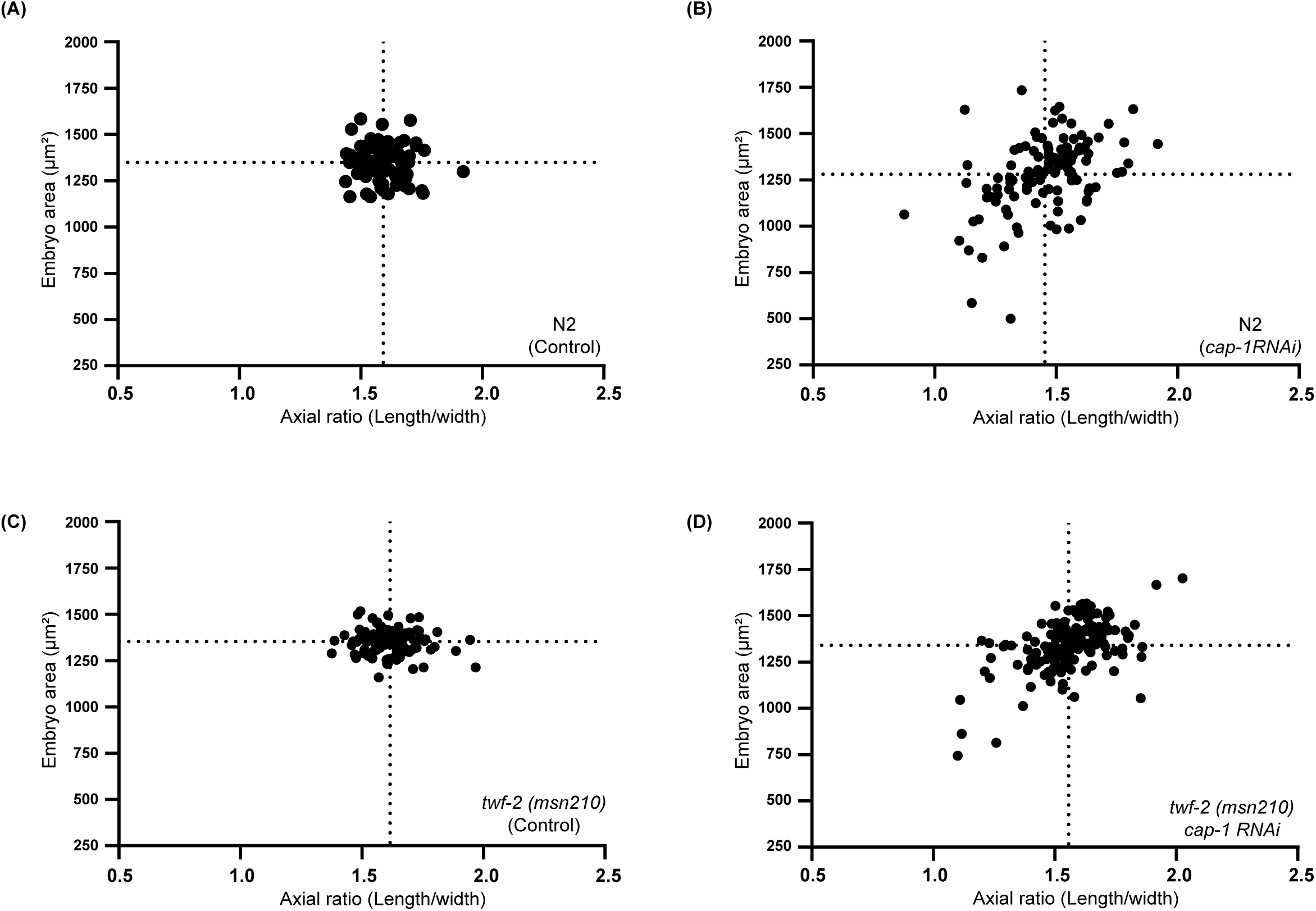
**Embryonic morphology analysis in cap-1(RNAi)** Correlation plots show embryo area versus axial ratio in wild-type(N2) in control and *cap-1(RNAi); twf-2 (msn210) mutant,* in control and *cap-1(RNAi)*. Dotted lines indicate the mean values. n≥87 embryos per genotype.

**Supplementary Figure 4.**
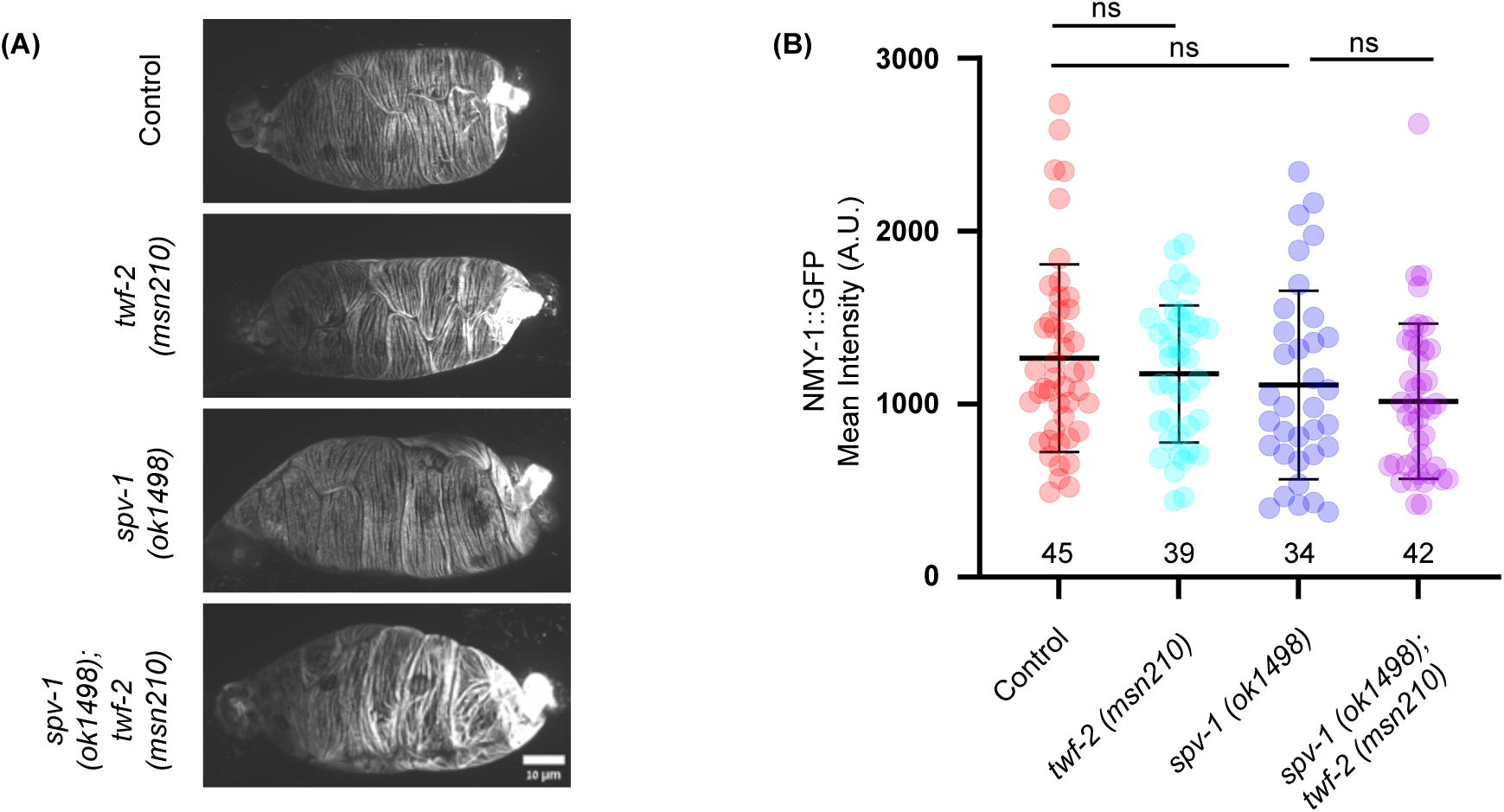
**Effects of *twf-2 and spv-1* mutations on endogenous NMY-1::GFP intensity.** (A) Representative maximum intensity projections for confocal images for endogenous NMY-1::GFP strain intensity in control, *twf-2 (msn210), spv-1 (ok1498),* and *twf-2(msn210);spv-1(ok1498)* double mutant strains. Scale bar: 10μm (B) Quantification of NMY-1::GFP fluorescence intensity in spermatheca. Horizontal and vertical bars represent Mean± SD. n≥34 spermathecae were scored for each genotype.

